# EP400NL is required for cMyc-mediated PD-L1 gene activation by forming a transcriptional coactivator complex

**DOI:** 10.1101/2021.05.30.446361

**Authors:** Zidong Li, Helen L Fitzsimons, Tracy K Hale, Jeong Hyeon Park

## Abstract

EP400 is an ATP-dependent chromatin remodeling enzyme that has been implicated in DNA double-strand break repair and transcription regulation including Myc-dependent gene expression. We previously showed that the N-terminal domain of EP400 increases the efficacy of chemotherapeutic drugs against cancer cells. As the EP400 N-terminal-Like (EP400NL) gene resides next to the EP400 gene locus prompted us to investigate whether EP400NL also plays a similar role in epigenetic transcriptional regulation to the full-length EP400 protein. We found that EP400NL forms a human NuA4-like chromatin remodelling complex that lacks both the TIP60 histone acetyltransferase and EP400 ATPase. However, this EP400NL complex displays H2A.Z deposition activity on a chromatin template comparable to the human NuA4 complex, suggesting another associated ATPase such as BRG1 or RuvBL1/RuvBL2 catalyses the reaction. We also demonstrated that the transcriptional coactivator function of EP400NL is required for cMyc and IFNγ-mediated PD-L1 gene activation. Collectively, our studies show that EP400NL plays a role as a transcription coactivator for cMyc-mediated gene expression and provides a potential target to modulate PD-L1 expression in cancer immunotherapy.

## INTRODUCTION

The E1A binding protein, EP400 ATPase is one of the subunits of human NuA4 (hNuA4) histone acetyltransferase (HAT) complex that regulates transcription through chromatin remodelling of target genes (1,2). In addition to EP400, the human NuA4 (hNuA4) complex consists of over 16 subunits including the phosphatidylinositol 3-kinase family protein kinase homolog TRRAP (Transformation/transcription domain-associated protein), TIP60 histone acetyltransferase, and bromodomain-containing BRD8 (2). The EP400 containing hNuA4 complex catalyses an ATP-dependent H2A.Z deposition activity to regulate gene expression and DNA double-strand break repair (3,4). The EP400 complex also promotes H3.3 histone variant deposition at certain target gene loci, suggesting that EP400-mediated double histone variant (H2A.Z/H3.3) deposition could play a key role in the epigenetic regulation of these genes (5,6). In connection, cMyc, a global transcription amplifier, is known to recruit the hNuA4 complex, and its interactions with TRRAP and EP400 are required for cMyc mediated gene activation (7,8).

One of the genes regulated by cMyc is programmed death ligand-1 (PD-L1), which is required for immune checkpoint control. The ligand-receptor interaction between PD-L1 and PD-1 has been extensively studied as a target of cancer immunotherapy ever since it was implicated in the suppression of cytotoxic T lymphocytes and immune tolerance, ultimately leading to tumor immune escape (9). cMyc has been demonstrated to bind to the PD-L1 gene promoter and induce expression, however, the molecular mechanisms through which this occurs are not well understood (10,11). In addition to Myc mediated PD-L1 expression, the pro-inflammatory cytokine interferon-gamma (IFNγ) also induces PD-L1 expression in a variety of tumors including melanoma, non-small cell lung cancer (NSCLC), and renal cell carcinoma (12-14). This phenomenon has been referred to as innate immune resistance which is a survival strategy of tumor cells to escape immune surveillance (15-17). IFNγ activates the IFNγ receptor, which further activates various signaling pathways such as PI3K/AKT/mTOR and JAK-STAT (18,19). The activation of these pathways can ultimately recruit downstream transcription factors such as IRF1, HIF-1α/2α, STAT1 dimers, STAT3, Myc, and NF-κB to the gamma interferon activation site (GAS) elements or other regulatory sites within the promoter region of PD-L1 to induce expression (18-24).

We previously showed that the N-terminal domain of EP400 (1-719 amino acids sequence) increases the efficacy of chemotherapeutic drugs against U2OS cancer cells that overexpress the N-terminal fragment of EP400 (25). Interestingly, we found that an EP400 N-terminal-Like (EP400NL) gene resides next to the EP400 gene on chromosome 12q24.33. The sequence alignment of EP400 and EP400NL shows that the N-terminal fragment of EP400 (131-490 amino acids) shares 92% sequence similarity over 84.5% of the EP400NL amino acids sequence. Despite no previous identification of a functional domain and its subsequent annotation as a pseudogene, EP400NL gene expression appears to be regulated by an independent promoter sequence, and the exon regions are highly conserved in many vertebrates (UCSC genome browser). Given their adjacent genetic location and sequence similarity, we hypothesized that EP400NL may have a regulatory function distinct from its homolog EP400 in the regulation of EP400-target genes. Here, we report that EP400NL forms a unique nuclear complex similar to the EP400 chromatin remodelling complex and serves as a transcriptional coactivator in cMyc and IFNγ-mediated PD-L1 transcriptional activation.

## MATERIAL AND METHODS

### Mammalian cell culture

FLP-In T-REX cells were maintained in Dulbecco’s Modified Eagle’s Medium (Gibco/Life Technologies) with 15 μg/ml blasticidin and 100 μg/ml Zeocin. FLP-In T-REX cells that stably express EP400NL were maintained in the same media with 15 μg/ml blasticidin and 30 μg/ml hygromycin. Unless specified, cells were maintained in Dulbecco’s Modified Eagle’s Medium (DMEM, Gibco/Life Technologies) supplemented with 5% fetal bovine serum (FBS) and 0.5% Penicillin-Streptomycin in a 5% CO2 incubator. For serum starvation, cells were grown for 48 h in the DMEM medium containing 1% FBS and then stimulated with 20% FBS or IFNγ (5 ng/ml and 20 ng/ml).

### Generation of EP400NL indel mutation by CRISPR/Cas9

Guide RNAs were designed based on a Toronto KnockOut (TKO) CRISPR Library for EP400NL targeted guide RNA (AGGTTGTGGCCAGAAAGCAC), luciferase targeted guide RNA (AACGCCTTGATTGACAAGGA), and scrambled guide RNA (AAACATGTATAACCCTGCGC). The guide RNAs were cloned into the enhanced specificity CRISPR/Cas9 plasmid (eSpCas9-LentiCRISPR, v2, Genscript) (26). HEK293T cells were seeded in a 10 cm plate at a density of 1 × 10^6^ cells per plate and used for lentiviral production. The lentiviral medium was subsequently concentrated using Lenti-X™ Concentrator (Takara Bio) and used to infect H1299 cells. The candidate cells were selected with 2 µg/ml puromycin for a week and the selected cells were analysed by High-resolution melting peaks analysis using real-time PCR (Light cycler 480 II, Roche).

### Subcellular fractionation

Cells that stably express EP4000NL were generated using the Flp-In™ T-REx™ System (Invitrogen). FLP-In T-REX cells were washed and scraped after tetracycline induction. Cells were pelleted and resuspended with hypotonic buffer (10 mM Tris pH 7.3, 1 mM KCl, 1.5 mM MgCl2, 0.01 M KCl) and the supernatant was removed after centrifugation. Hypotonic buffer was then added and homogenized with size B pestle of Dounce homogenizer. Cells were spun at 4000 rpm for 15 m and the supernatant was used as the cytosolic fraction. After the spin, the nuclear pellet was resuspended with low salt buffer (20 mM Tris pH 7.3, 12.5% Glycerol, 1.5 mM MgCl_2_, 0.2 mM EDTA, 20 mM KCl) and transferred for homogenization. High salt buffer (20 mM Tris pH 7.3, 12.5% Glycerol, 1.5 mM MgCl_2_, 0.2 mM EDTA, 1.2 M KCl) was added to the homogenized nuclear pellet at 4 °C and stirred for 30 m followed by 30 m centrifugation at 30,000 rpm. After taking the supernatant as the nuclear soluble fraction, the nuclear pellet was then resuspended and centrifuged at the same speed for nuclear pellet fraction.

### Purification and elution of EP400NL complex

Approximately 5 × 10^8^ cells of tetracycline-induced FLP-In T-REX cell line were harvested and prepared for a nuclear soluble fraction. The fraction was subjected to gel filtration chromatography using Bio-Gel A1.5 resin equilibrated with streptavidin binding buffer (10 mM Tris, pH 8.0, 150 mM NaCl, 2 mM EDTA pH 8.0, 0.1% NP40, and 10 mM 2-Mercaptoethanol). Tetracycline-induced EP400NL protein, tagged with both streptavidin-binding peptide (SBP) and calmodulin-binding peptide (CBP) was affinity-purified by using streptavidin-conjugated agarose beads (EZview™ Red Streptavidin Affinity Gel, Sigma-Aldrich). The initial protein peaks containing EP400NL were collected and pooled together after the size exclusion chromatography and incubated with streptavidin affinity beads at 4 °C overnight with constant rotation. The beads were collected and washed followed by biotin elution. Eluted protein samples were used as an input source for mass spectrometric analysis.

### H2A.Z deposition assay

A plasmid containing 19 repeats of 601-Widom nucleosome positioning sequences (pUC19/601×19) was used to assemble the chromatin substrate. Biotin labelling of the nucleosome assembly was conducted using the EZ-Link™ Psoralen-PEG3-Biotin kit (Thermo Scientific™) as described in the manufacturer’s protocol. hNuA4 complex was purified from Hela cell nuclear extracts expressing FLAG-TIP60. All FLAG-tagged proteins were purified with anti-FLAG® M2 Affinity Gel (Sigma-Aldrich) in a native condition using the FLAG peptide elution. The partially purified enzyme fractions were added to the reaction mixture containing 100 ng FLAG-H2A.Z/H2B dimer, 5 μg assembled chromatin, 1× protease inhibitor (Roche), and 1 mM ATP and incubated for 1 h at 30 °C. The chromatin was harvested by the streptavidin-conjugated agarose beads (EZview™ Red Streptavidin Affinity Gel, Sigma-Aldrich), and analysed by immunoblotting.

### Dual-luciferase reporter assays

HEK293TGal4-Luciferase cells were seeded in 24-well plates and transfected with CbF-Gal4Myc (20 ng), pRenilla-CMV (1ng), and plasmids expressing various cofactors (200 ng) using Effectene Transfection Reagent (QIAGEN). After 48 hours, firefly luciferase and renilla luciferase activities were measured using the Dual - Luciferase® Reporter Assay System (Promega) and the POLAR star Omega plate reader (BMG LABTECH). Triplicate experiments were conducted and averaged firefly luciferase activity was normalized to the corresponding Renilla activity.

### Chromatin immunoprecipitation-quantitative PCR (ChIP-qPCR) assays

Chromatin immunoprecipitation (ChIP) assays were performed according to the protocol obtained from Abcam high-sensitivity ChIP kit (ab185913, Abcam) with several modifications. HEK-293TGal4-Luciferase cells were treated with 1% formaldehyde for 5 min followed by incubation in 0.125 M glycine for 5 min at room temperature. Cells were washed, lysed, and sonicated to a size of approximately 500 bp. The 20 µg of sonicated DNA per reaction was immunoprecipitated overnight with antibodies against FLAG (A8592, Sigma-Aldrich), CBP (sc-33000, Santa Cruz Biotechnology), BRG1 (# 710894, Invitrogen), RuvBL1, RuvBL2 (kind gifts from Dr. Cole, Dartmouth Medical School) and IgG which was included in the Abcam high-sensitivity Chip kit (ab185913). Immune complexes were collected using protein A/G PLUS-Agarose beads (sc-2003, Santa Cruz Biotechnology), washed once with LiCl_2_ buffer, and eluted in 50 µl fresh elution buffer (1% SDS (v/v), 50 mM Tris, pH 7.0, 10 mM EDTA). The DNA was reverse cross-linked and purified using phenol-chloroform and ethanol. Protein was removed by incubating with proteinase K at 65 °C for 3 h. The purified DNA was quantified using the Luna® Universal qPCR Master Mix (NEB) for the specific targeted regions of the Gal4 DNA binding site and PD-L1 promoter region respectively. The antibodies and primer information (27,28) are summarised in supplementary tables 1 and 2.

### Confocal microscopy

Cells were seeded in a 6 well plate at a density of 1.5×10^5^/ml together with two UV sterilised coverslips in each well for overnight culture. Cells were washed with PBS (phosphate-buffered saline) and fixed with 2% paraformaldehyde in PBS for 15 m at room temperature followed by the permeabilization with 0.2% Triton X-100 in PBS for 5 m with a gentle mix every few minutes and subsequently blocked for 60 m in PBS containing 5% bovine serum albumin (BSA) and 0.05% Tween-20. Coverslips were overlaid with primary antibodies diluted in blocking buffer and incubated overnight at 4 °C. The primary antibody solution was removed in the next day, followed by incubating for 60 m at room temperature with the secondary antibodies diluted in blocking buffer. Coverslips were washed and fixed in 2% paraformaldehyde in PBS for 15 m at room temperature followed by a final wash with distilled H_2_O. The slide was examined by Zeiss LSM900 confocal laser scanning microscope.

### Statistical analysis

Statistical analysis was performed using GraphPad Prism software version 8.0 (GraphPad Software Inc., San Diego, CA, USA). Data were compared with either an ordinary one-way ANOVA statistical test or a two-way ANOVA statistical test depend on the experimental design and followed by either one of the following tests which are Dunnett’s multiple comparisons, Tukey’s multiple comparisons, or Sidak’s multiple comparisons based on their individual data collection methods. P values less than 0.05 were considered statistically significant. (N.S. P > 0.05, * P ≤ 0.05, ** P ≤ 0.01, *** P ≤ 0.001, **** P ≤ 0.0001). All data shown were determined from independent experiments and presented as the mean ± SD.

## RESULTS

### EP400NL serves as a positive coactivator in GAL4Myc-mediated transcriptional activation

The sequence similarity of EP400NL to the N-terminal domain of EP400 prompted us to investigate a potential coactivator function of EP400NL. Given that the EP400 chromatin remodelling complex has been implicated in Myc-dependent transactivation (7,29), the activation of the luciferase reporter (30,31) by the GAL4Myc transcription activator was analysed in the presence of EP400NL in 293TGal4-Luciferase cells (Figure 1A). As a control, previously known coactivators of hNuA4 complex (TRRAP, EP400, and their defective mutants) were included to compare with EP400NL (Figure 1A). Consistent with the role as a positive coactivator, TRRAP stimulated luciferase expression up to two-fold, whereas a C-terminal deletion mutant of TRAAP failed to promote similar gene activation (32-34). Although modest, EP400 also showed consistent stimulation of reporter gene expression whereas the ATPase-deficient EP400 did not. Interestingly, EP400NL induced reporter activity up to 2.2-fold, suggesting that EP400NL activates transcription at a level comparable to that of TRRAP. To examine a dose-dependent effect of EP400NL, luciferase expression was examined in the presence of increasing ectopic expression of EP400NL in 293TGal4-Luciferase cells (Figure 1B, top panel). In the presence of a similar GAL4Myc level, the reporter gene expression showed a dose-dependent increase in expression of up to three-fold in the presence of increasing amounts of EP400NL (Figure 1B, bottom panel).

**Figure 1.**
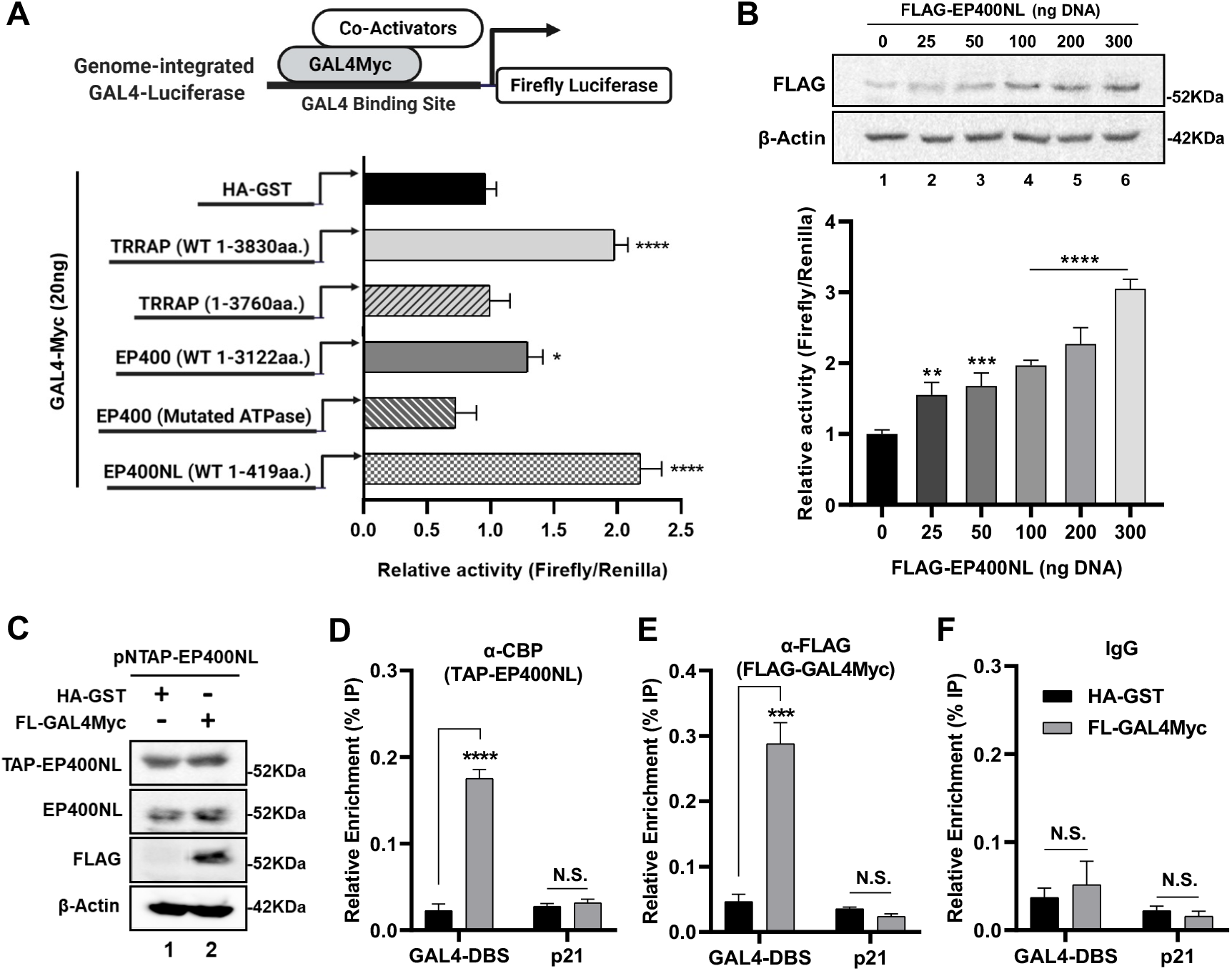
EP400NL serves as a positive coactivator in GAL4Myc-mediated transcriptional activation. (A) Firefly luciferase reporter assays in 293TGal4-Luciferase cells. The cells were transfected with each construct expressing either HA-GST (control), wild type TRRAP (1-3830aa.), TRRAP C-terminal deletion mutant (1-3760aa.), wild type EP400 (1-3122aa.), ATPase mutated EP400, and wild type EP400NL (1-419aa.) together with a plasmid expressing GAL4Myc, respectively. After normalised by the renilla luciferase activity, the firefly luciferase activity from the control cells (HA-GST) set to one and the relative values for other samples are calculated [Ordinary one-way ANOVA, *F* _*(5,12)*_ = 55.96, *p* < 0.0001; Dunnett’s post-hoc test, *\*p* < 0.05, *\*\*\*\*p* < 0.0001]. (B) Titration of EP400NL expression. Transient transfection of CbF-EP400NL (0 ng, 25 ng, 50 ng, 100 ng, 200 ng, 300 ng) into HEK293TGal4-Luciferase cells was conducted for the dual-luciferase reporter assay [Ordinary one-way ANOVA, *F* _*(5,12)*_ = 61.11, *p* < 0.0001; Dunnett’s post-hoc test, *\*\*p* < 0.01, *\*\*\*p* < 0.001, *\*\*\*\*p* < 0.0001]. Protein expression of EP400NL was confirmed by anti-FLAG immunoblot (top panel) and the relative firefly luciferase activities were plotted against the amount of DNA used in the transfection (bottom panel). (C) 293TGal4-Luciferase cells were co-transfected with plasmids expressing either HA-GST (control) or FLAG-GAL4Myc together with TAP-EP400NL respectively. Protein expression was confirmed by immunoblotting. (D-F) ChIP analyses were performed using anti-CBP for TAP-tagged EP400NL (D), anti-FLAG for GAL4Myc (E), and IgG as a negative control (F). The ChIP data are plotted for relative enrichment of input chromatin from each ChIP reaction over the GAL4 DNA binding site (GAL4-DBS) and p21 distal exon region (p21). [Two-way ANOVA, *F* _*(1,4)*_ = 225.7, *p* = 0.0001 (CBP), *F* _*(1,4)*_ = 107.0, *p* = 0.0005 (FLAG); Sidak’s post-hoc test, *\*\*\*p* < 0.001, *\*\*\*\*p* < 0.0001].

To demonstrate activator-dependent recruitment of EP400NL on target genes, chromatin immunoprecipitation (ChIP) was performed with 293TGal4-Luciferase cells in the presence or absence of ectopic FLAG-GAL4Myc and TAP-tagged EP400NL. Expression was confirmed by immunoblot analysis (Figure 1C). The enrichment of EP400NL at the GAL4 binding site (GAL4-DBS) was significantly increased by the presence of GAL4Myc by approximately 8-fold, indicating that the recruitment of EP400NL is dependent on GAL4Myc (Figures 1D, 1E, and 1F). The distal exon region of p21 was used as a control to establish the specificity of the experiment where neither of the GAL4Myc nor EP400NL was enriched in that region. Taken together, these results suggest that EP400NL promotes reporter gene expression by the specific recruitment onto the promoter in a DNA-binding transcription factor-dependent manner.

### EP400NL forms a unique nuclear complex similar to the EP400 chromatin remodelling complex

To identify EP400NL-associated proteins, the Flp-In™ T-REx™ cell line was used to inducibly express TAP-tagged EP400NL in a tetracycline-dependent manner. Inducible expression of the TAP-tagged EP400NL and the putative endogenous EP400NL were confirmed by immunoblotting (Figure 2A). The cytosolic (CT), soluble nuclear (SN), and insoluble nuclear fractions (IN) were analyzed by immunoblotting to identify the subcellular localization of EP400NL, which was found to localize predominantly to the soluble nuclear fraction (Figure 2B, left panel). PPM1B, MED30, and Lamin A/C are used as a marker of cytosol, nucleoplasm, and an insoluble component of the nuclear pellet, respectively (35-37). The localization of tetracycline-induced TAP-EP400NL in the nucleus was further verified by confocal microscopy that shows immunochemical staining of TAP-EP400NL and DAPI stained nuclei together (Figure 2B, right panel).

**Figure 2.**
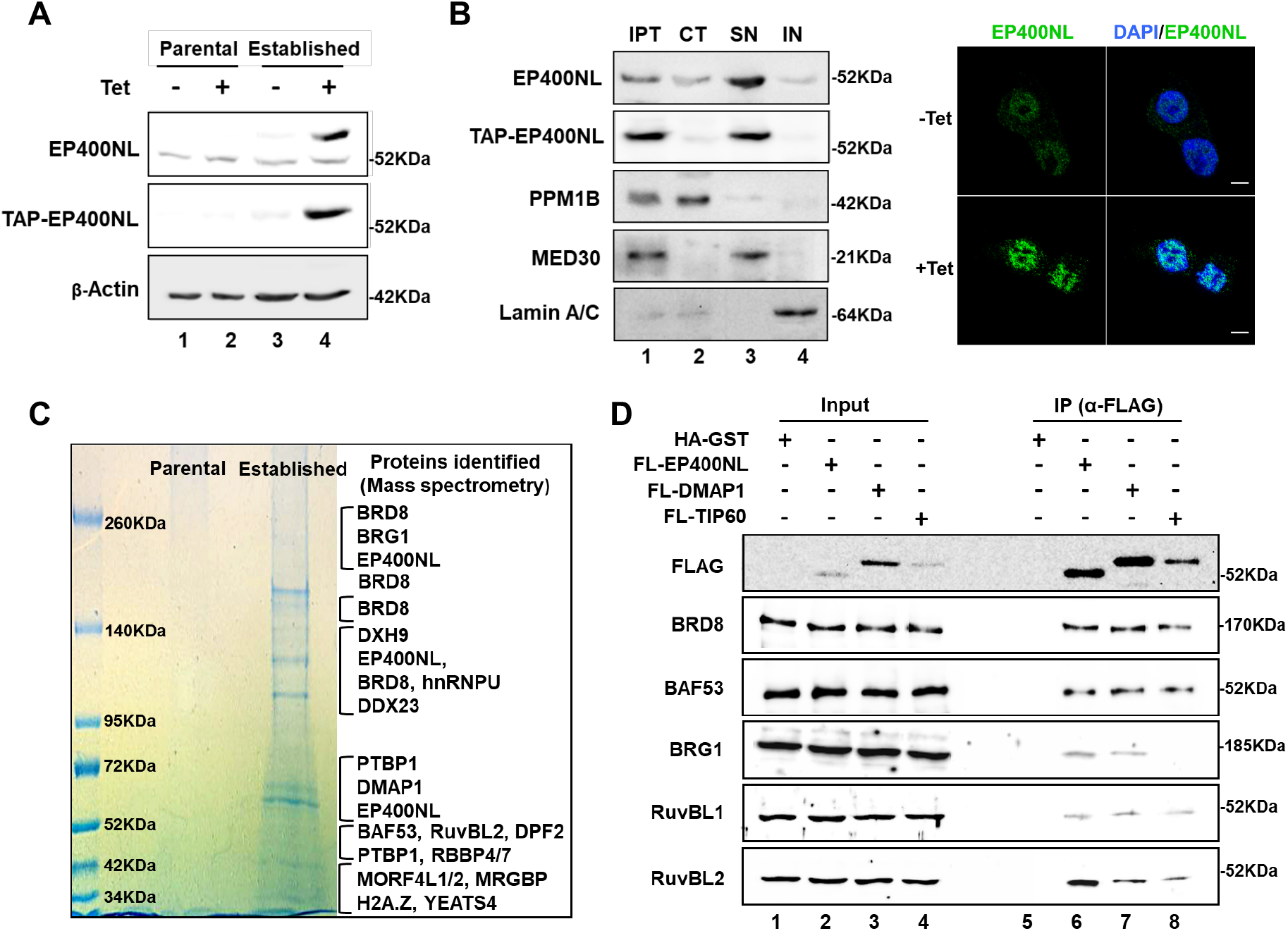
EP400NL forms a nuclear complex similar to the EP400 chromatin remodelling complex. (A) Immunoblot analysis of Flp-In™ T-REx™ cell line stably expressing tetracycline-inducible EP400NL. TAP-tagged EP400NL (approximately 60 KDa in molecular mass) and putative endogenous EP400NL (around 52kDa in molecular mass) are shown by anti-EP400NL and anti-CBP tag immunoblots. (B) Cellular fractionation of EP400NL. Input control (IPT), cytosolic fraction (CT), soluble nuclear fraction (SN), and insoluble nuclear pellet (IN) were analysed by immunoblotting (left panel). Subcellular localization of TAP-EP400NL was analysed by immunocytochemical analysis (right panel). TAP-EP400NL inducible Flp-In™ T-REx™ cell line was treated with or without tetracycline. Scale bar denotes 10 µm. (C) Colloidal Coomassie Blue G250 staining of EP400NL nuclear complex. Protein samples were purified from Flp-In™ T-REx™ cell lines either expressing TAP-EP400NL or parental control, respectively. The gel pieces after SDS-PAGE and protein staining were analysed by mass spectrometry. Identified proteins are indicated on the right side of the stained gel. (D) Interaction of EP400NL with the candidate proteins by co-immunoprecipitation assay. HEK293T cells were transiently transfected with plasmids expressing HA-GST, FLAG-EP400NL, FLAG-DMAP1, or FLAG-TIP60 and the cell lysates were immunoprecipitated by anti-FLAG antibody. The immunoprecipitates were analysed by immunoblots using anti-FLAG, anti-BRD8, anti-BAF53, anti-BRG1, anti-RuvBL1, and anti-RuvBL2 antibodies.

To identify EP400NL-associated proteins, the TAP-tagged EP400NL complex was isolated through a combination of size exclusion chromatography and SBP affinity tag purification from the soluble nuclear fraction of the induced cell line. The purified protein samples derived from parental and established FLP-In T-REX cells were resolved by SDS-PAGE and identified by mass spectrometry (Figure 2C). Not surprisingly, most of the identified proteins were integral components of hNuA4 chromatin remodelling complexes including BRD8, BAF53, DMAP1, RuvBL2 (TIP48), MRGBP, and YEATS (2,38).

To verify the mass spectrometric identification, three FLAG-tagged proteins (EP400NL, DMAP1, and TIP60) were transiently expressed in HEK293T cells and partially purified by anti-FLAG M2 agarose beads (Figure 2D). Consistent with the results obtained from the mass spectrometry, EP400NL co-immunoprecipitated with BRG1, BRD8, BAF53, and RuvBL2. RuvBL1 is known to form a hetero-oligomeric complex with RuvBL2 in most human nuclear complexes (39). Although RuvBL1 was not identified by the mass spectrometry it also appears to be a potential interacting partner with EP400NL. Interestingly, BRG1, a core ATPase subunit of the human SWI/SNF chromatin remodelling complexes, was identified as a component of the EP400NL complex by the mass spectrometry and further confirmed by co-immunoprecipitation assay (Figure 2D, α-BRG1 blot, lane 6). BRG1 has not been reported as an interacting protein with the hNuA4 complex (1,2,38), however, it seems to associate with both EP400NL and DMAP1 (Figure 2D, α-BRG1 blot, lanes 6 and 7).

### The EP400NL nuclear complex catalyses an H2A.Z deposition to chromatin *in vitro*

The EP400NL nuclear complex appears to be a distinct complex from the hNuA4 complex because it does not appear to contain TRRAP, EP400, or TIP60 HAT. Further, a partially purified EP400NL complex did not show any HAT activity with chromatin as a substrate (data not shown).

The identification of H2A.Z as an interacting protein with EP400NL by mass spectrometry led us to investigate whether this complex mediates ATP-dependent H2A.Z deposition as previously shown by the EP400 complex (4,40). FLAG-tagged H2A.Z/H2B histones were used as a substrate to investigate if a partially purified FLAG-tagged EP400NL complex can catalyse H2A.Z deposition into chromatin (Figure 3A). As a control, the hNuA4 complex was purified from HeLa nuclear extract using FLAG-tagged TIP60, and the presence of each core protein (RuvBL1, RuvBL2, BRG1, and BAF53) was confirmed by immunoblotting (Figure 3B). Consistent with the mass spectrometry data, BRG1 was present in the EP400NL complex but not the hNuA4 complex. Plasmids containing the 601-Widom repeats (pUC19/601×19) were biotinylated and used to generate synthetic chromatin with recombinant canonical histones. Synthetic chromatin assemblies were validated by MNase digestion that showed a 200bp spaced ladder (Figure 3C). The EP400NL complex displayed an increase in FLAG-H2A.Z deposition activity over background, as did the hNuA4 complex in the presence of ATP (Figure 3D).

**Figure 3.**
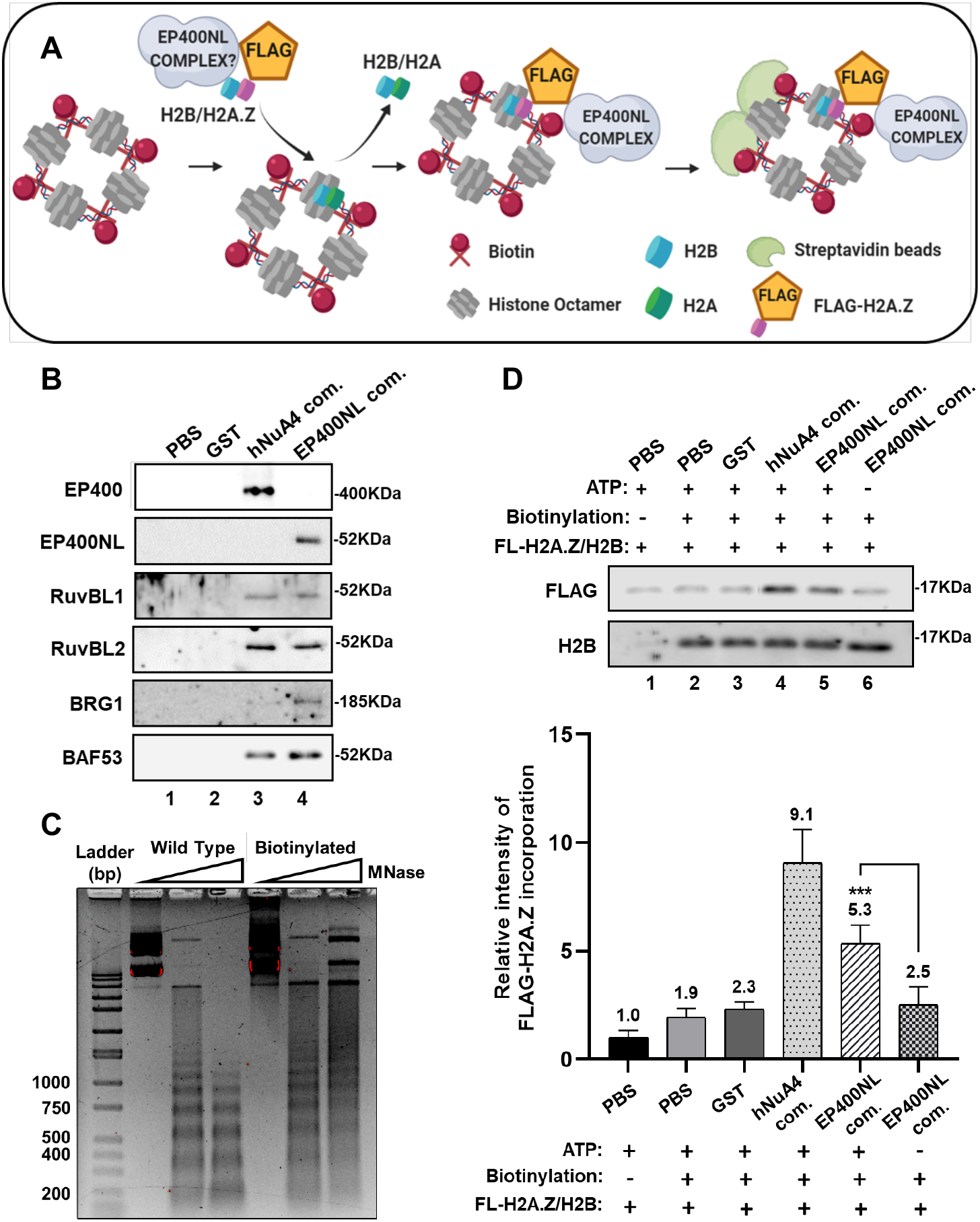
EP400NL nuclear complex catalyses H2A.Z deposition activity to the chromatin *in vitro*. (A) Schematic illustration of H2A.Z deposition assay. (B) Protein preparations used in the H2A.Z deposition assay were analyzed by immunoblots for the core components of the protein complexes including EP400, EP400NL, RuvBL1, RuvBL2, BRG1, and BAF53. (C) MNase digestion assay of the assembled chromatin. Chromatin substrates assembled with either biotin-labeled pUC19/601×19 or unlabelled plasmid DNA were digested by two different concentrations of MNase and analysed by 1% agarose gel electrophoresis. (D) Detection of FLAG-H2A.Z incorporation to the chromatin substrate by immunoblotting. The reactions were resolved by SDS-PAGE and H2A.Z deposition activities were analyzed by anti-FLAG immunoblot (top panel). The relative intensity of FLAG-H2A.Z was initially normalized by the H2B signals and the normalized value from the negative control (PBS and non-biotinylated chromatin) sets to one and the relative activities from other samples were plotted (bottom panel) [Ordinary one-way ANOVA, *F* _*(5,24)*_ = 65.75, *p* < 0.0001; post-hoc Tukey’s HSD, *\*\*\*p* < 0.001].

### Functional analysis of EP400NL in GAL4Myc-mediated transcriptional regulation

No functional domains within EP400NL have been identified to date, however, a bioinformatic prediction tool InterPro which is a powerful integrated database of protein families, domains, and functional sites, predicts seven potential functional regions within EP400NL (Figure 4A). To investigate the importance of these regions in the interactions with BAF53, BRD8, and BRG1, full-length EP400NL and four deletion mutants of EP400NL (Δ1-50, Δ141-156, Δ246-260, Δ360-419) were generated and transiently expressed in HEK293T cells (Figure 4B). The mutants were expressed at relatively higher levels to wild-type EP400NL, despite all of them retained the ability to interact with BRD8 and BRG1, two deletion mutants of EP400NL (Δ246-260, Δ360-419) showed a weaker interaction with BAF53 (Figure 4B).

**Figure 4.**
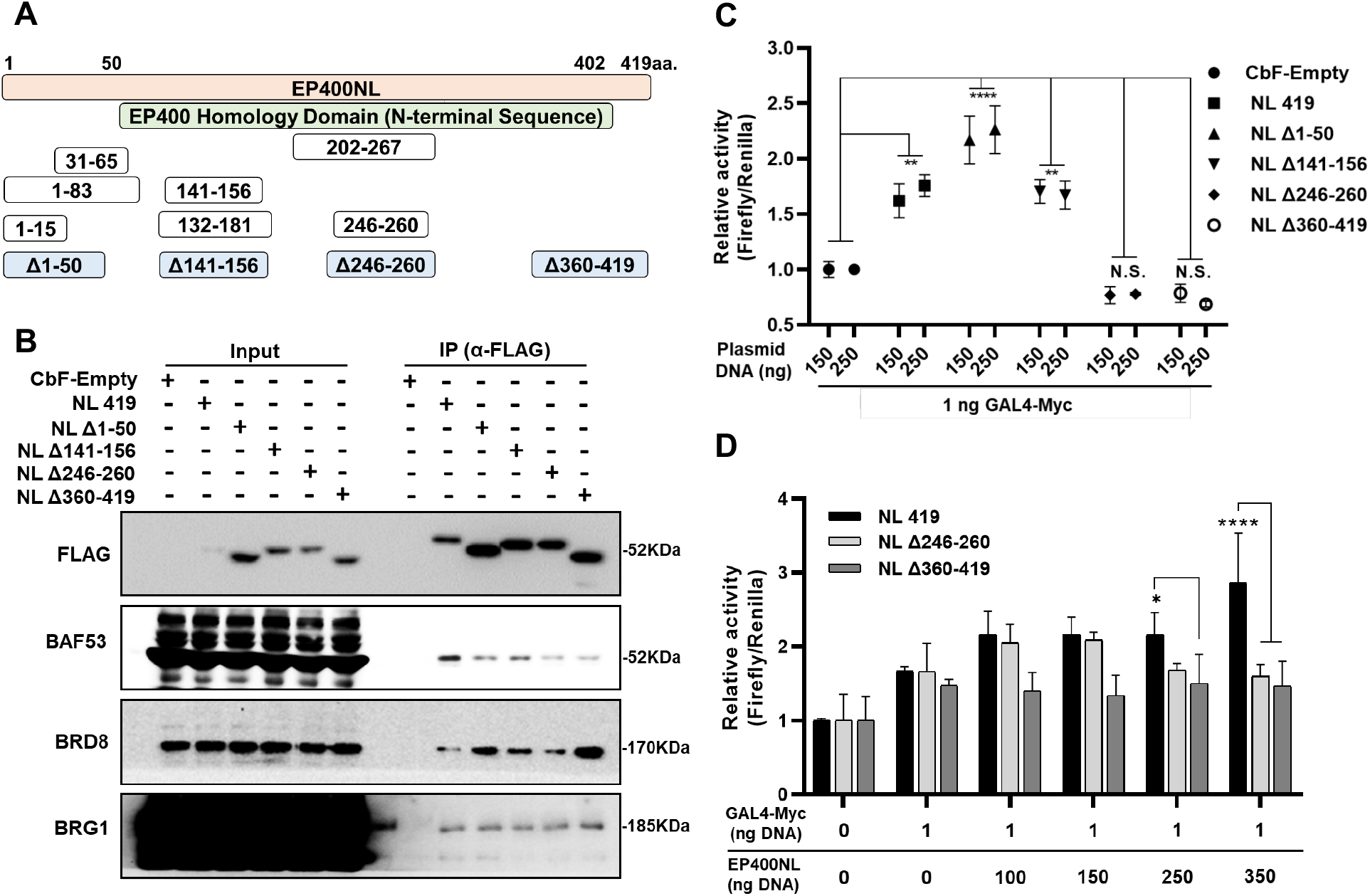
Functional analysis of EP400NL and its mutants in GAL4Myc-mediated transcriptional activation. (A) Schematic presentation of four deletion mutants of EP400NL (Δ1-50, Δ141-156, Δ246-260, Δ360-419). White boxes represent the seven predicted functional sites by the bioinformatic tool (InterPro). (B) Co-immunoprecipitation assays of EP400NL wild type and deletion mutants. HEK293T cells were transiently transfected with plasmids expressing FLAG-tagged EP400NL or its mutants for partial immunopurification with anti-FLAG antibody. (C) Firefly luciferase reporter assays in 293TGal4-Luciferase cells. The cells were transfected with either 150 ng or 250 ng of plasmid DNA expressing wild-type EP400NL or its deletion mutants together with 1 ng of Gal4Myc expression plasmid. After normalised by the renilla luciferase activity, the firefly luciferase activity from the control cells (GAL4Myc expression + empty vector) set to 1 and the relative values of other samples are calculated [Two-way ANOVA, *F* _*(5,24)*_ = 50.28, *p* < 0.0001; post-hoc Tukey’s HSD, *\*\*p* < 0.01, *\*\*\*\*p* < 0.0001]. (D) Similar luciferase assays were performed for two deletion mutants (Δ246-260, Δ360-419) by increasing the amount of plasmid DNA in the transfection reactions. After normalization to renilla luciferase, firefly luciferase activity from the control cells (no GAL4Myc expression) was set to 1 and the relative values for other samples are calculated [Two-way ANOVA, *F* _*(2,36)*_ = 21.11, *p* < 0.0001; post-hoc Tukey’s HSD, *\*p* < 0.05, *\*\*\*\*p* < 0.0001].

To examine whether the mutants also retained transcriptional coactivator activity, they were tested for their ability to co-activate GAL4Myc-mediated luciferase reporter expression. Compared to the deletion of Δ1-50 which showed a modest upregulated coactivator activity, the deletion of either Δ246-260 or Δ360-419 resulted in the loss of coactivator activity compared to full-length EP400NL (Figure 4C). To further confirm the loss of coactivator function, plasmid DNA expressing full-length EP400NL or either mutant was titrated to examine a possible dominant-negative effect. As the amount of plasmid DNA transfected increased, the C-terminal deletion mutant, Δ360-419 shows a significant decrease compared to the full-length EP400NL but failed to further decrease below the basal level of GAL4Myc-mediated gene expression (Figure 4D). Taken together, these data show that regions within amino acids 246-260 and 360-419 of EP400NL are essential for transcriptional coactivation.

### EP400NL is targeted to the PD-L1 promoter in a cMyc-dependent manner

Given that EP400NL was recruited to the Gal4 DNA binding site of the promoter by the Myc transactivation domain (Figure 1D), possible involvement of EP400NL in the Myc-mediated PD-L1 gene regulation was explored.

To investigate whether EP400NL is recruited to the PD-L1 promoter in a Myc-dependent manner, ChIP was performed following induction of TAP-EP400NL expression in Flp-In™ T-REx™ cells. Cells were serum-starved to maintain a low growth rate with basal levels of Myc, then treated with high serum to promote cell growth. Despite the serum stimulation, no significant increase in cMyc enrichment was detected at the PD-L1 promoter (Figure 5A, cMyc panel), however, TAP-EP400NL induction enhanced cMyc enrichment upon serum-stimulation. Two potential ATPases (BRG1 and RuvBL2) (41,42) that were identified as the associated proteins with EP400NL were also examined for enrichment at the PD-L1 promoter. BRG1 and RuvBL2 showed a similar recruitment pattern in which both serum stimulation and tetracycline induction of EP400NL resulted in a significantly higher enrichment than serum stimulation or induction of EP400NL individually (Figure 5A, strapped bars versus dark or light grey bars). These data indicate that cellular signals induced by serum stimulation are a prerequisite for the enrichment of BRG1 and TIP48 ATPases near the Myc binding site of the PD-L1 gene, which can be significantly enhanced by the upregulated EP400NL.

**Figure 5.**
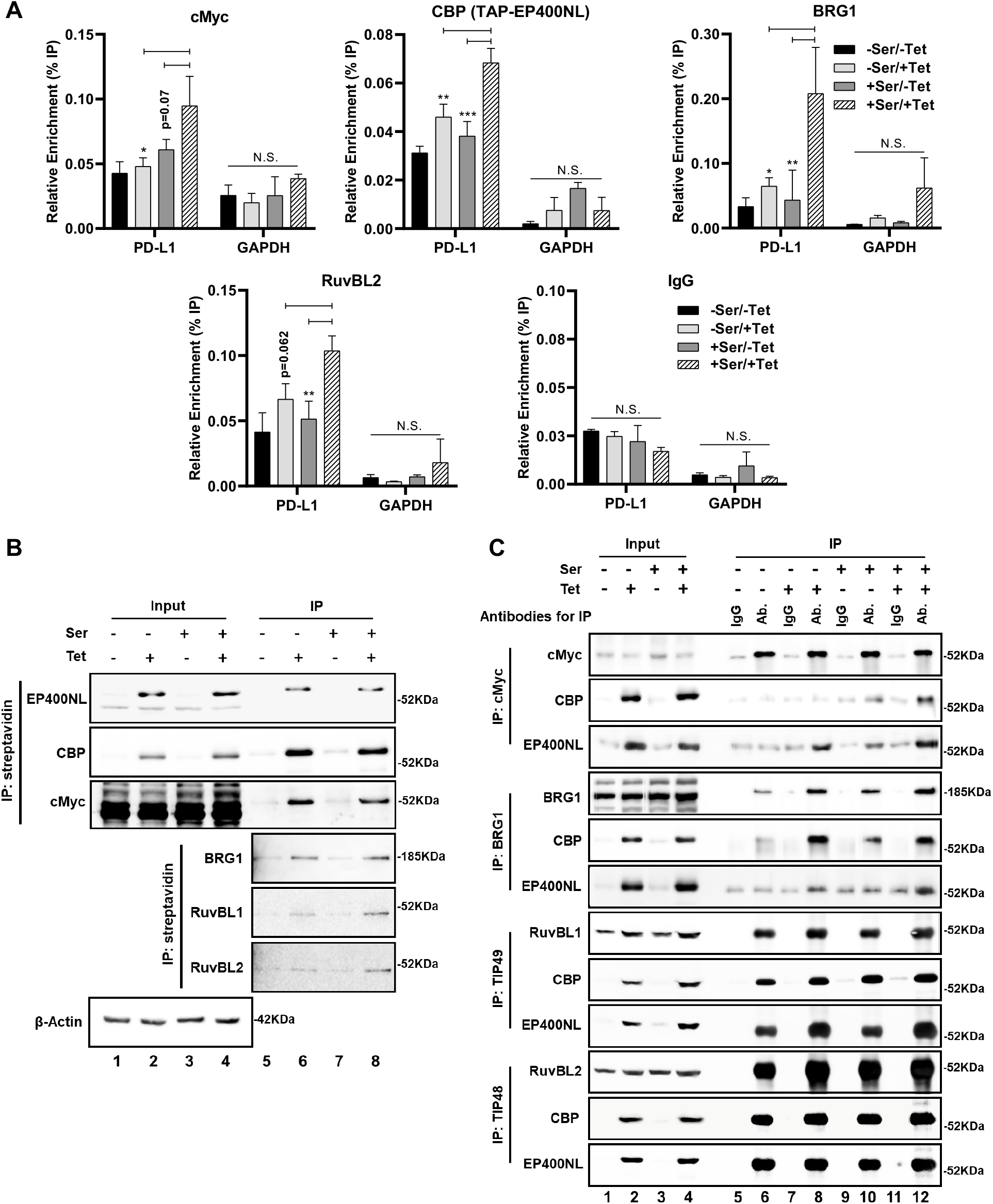
EP400NL is recruited at the PD-L1 promoter in a cMyc dependent manner. (A) Enrichment of cMyc, EP400NL, BRG1, and RuvBL2 at the PD-L1 promoter of Flp-In™ T-REx™ cell line stably expressing tetracycline-inducible EP400NL. The purified DNA after ChIP reactions was analyzed by qPCR over the regions of PD-L1 promoter (Myc binding site) or GAPDH promoter. Four combinations of the experimental conditions (the presence and absence of serum and tetracycline: ±Ser/±Tet) were used in the ChIP analyses [Two-way ANOVA, *F* _*(1,8)*_ = 36.01, *p* = 0.0003 (cMyc), *F*_*(1,8)*_ = 263.7, *p* < 0.0001 (CBP), *F* _*(1,8)*_ = 13.57, *p* = 0.0062 (BRG1), *F* _*(1,8)*_ = 103.6, *p* <0.0001 (RuvBL2); post-hoc Tukey’s HSD, *\*p* < 0.05, *\*\*p* < 0.01, *\*\*\*p* < 0.001]. (B) Co-immunoprecipitation experiments for the protein-protein interaction of EP400NL, cMyc, BRG1, RuvBL1, and RuvBL2. EP400NL protein complexes were immunoprecipitated by streptavidin beads followed by the validation of both anti-EP400NL and anti-CBP immunoblots. The co-immunoprecipitations of cMyc, BRG1, RuvBL1, and RuvBL2 were confirmed by the immunoblots. (C) Reverse co-immunoprecipitation experiments were performed with the same cell lysates in (B). Immunoprecipitates by anti-cMyc, anti-BRG1, anti-RuvBL1, and anti-RuvBL2 antibodies (Ab.) were examined by both anti-CBP and anti-EP400NL immunoblots. IgG immunoprecipitation serves as a negative control.

To examine if the interaction of cMyc with EP400NL, BRG1, RuvBL1, and RuvBL2 occurs under physiological conditions, a serum stimulation-based experiment was conducted using the stable Flp-In™ T-REx™ cell line expressing tetracycline-inducible TAP-EP400NL. Following tetracycline induction and serum stimulation, TAP-EP400NL was immunoprecipitated by streptavidin beads (Figure 5B) and the presence of endogenous cMyc, BRG1, RuvBL1, and RuvBL2 was detected in the immunoprecipitates (Figure 5B, lanes 6 and 8). Reverse immunoprecipitation using anti-cMyc, anti-BRG1, anti-RuvBL1, and anti-RuvBL2 antibodies also resulted in the immunoprecipitation of TAP-EP400NL (Figure 5C, lanes 6, 8, 10, and 12). These results further support a notion that cMyc directly interacts with the EP400NL nuclear complexes containing BRG1, RuvBL1, and RuvBL2 and those interactions can be further stabilised by EP400NL.

### EP400NL serves as a coactivator for both Myc and IFNγ-mediated induction of PD-L1 expression

To elucidate the role of EP400NL in regulating PD-L1 expression, the inducible Flp-In™ T-REx™ cell line was used to determine how the upregulation of TAP-EP400NL expression affects Myc-mediated PD-L1 expression. PD-L1 mRNA expression level increased up to approximately 2-fold by either serum stimulation or by tetracycline induction and the mRNA expression was further increased up to 3.2-fold in the presence of both serum stimulation and tetracycline induction (Figure 6A, strapped bar). Consistent with the effect on PD-L1 mRNA level, PD-L1 protein expression in the serum-stimulated and tetracycline induced cells is increased up to 2-fold compared to the untreated cells (Figure 6B, anti-PD-L1 immunoblot).

**Figure 6.**
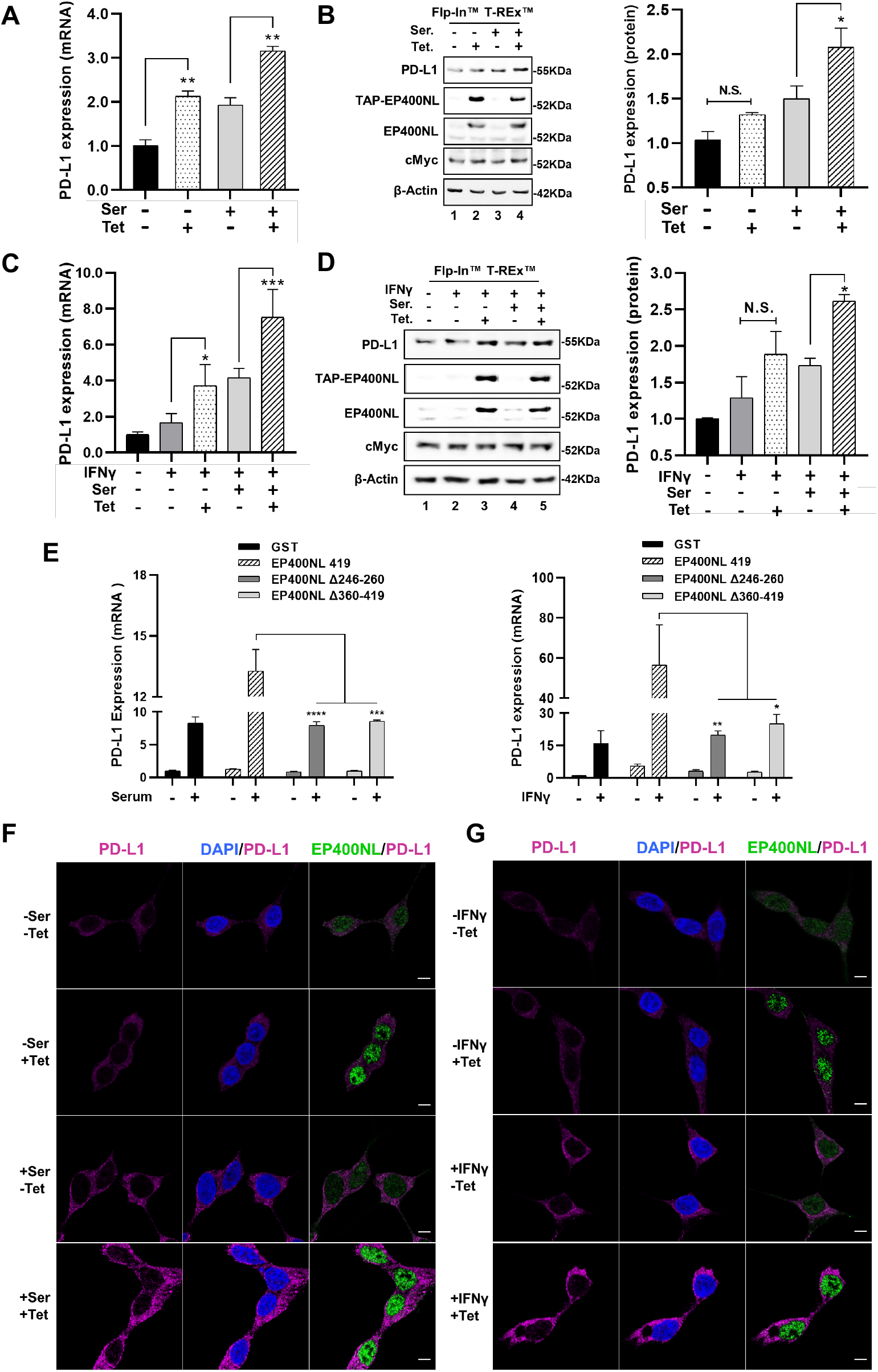
EP400NL serves as a coactivator for both Myc and IFNγ mediated PD-L1 gene activations. (A) PD-L1 mRNA expression levels in response to serum stimulation and tetracycline induction. Flp-In™ T-REx™ cells were treated under four different conditions (±Ser/±Tet) and the total RNA of each treatment was analysed by RT-qPCR assays [Ordinary one-way ANOVA, *F* _*(3,4)*_ = 85.42, *p* = 0.0004; post-hoc Tukey’s HSD, *\*\*p* < 0.01]. (B) Protein expression levels of endogenous PD-L1 in four different conditions (±Ser./±Tet.) were examined by immunoblots. The relative intensities of PD-L1 protein levels are quantified and plotted at the right panel [Ordinary one-way ANOVA, *F* _*(3,4)*_ = 20.32, *p* = 0.0070; post-hoc Tukey’s HSD, *\*p* < 0.05]. (C) PD-L1 mRNA expression level in response to a combination of serum stimulation, tetracycline induction, and IFNγ sensitization [Ordinary one-way ANOVA, *F* _*(4,15)*_ = 30.42, *p* < 0.007; post-hoc Tukey’s HSD, *\*p* < 0.05, *\*\*\*p* < 0.001]. (D) Protein expression levels of endogenous PD-L1 in five different conditions (±IFNγ/±Ser/±Tet) were examined by immunoblots. The relative intensities of PD-L1 protein levels are quantified and plotted at the right panel [Ordinary one-way ANOVA, *F* _*(4,5)*_ = 18.81, *p* = 0.0032; post-hoc Tukey’s HSD, *\*p* < 0.05]. (E) EP400NL mutants lose the coactivator function for both Myc- and IFNγ-mediated PD-L1 expression. H1299 cells were transiently transfected with plasmids expressing either wild type EP400NL or two deletion mutants (Δ246-260 and Δ360-419) and PD-L1 mRNA levels induced by serum stimulation (left panel) or IFNγ sensitization (right panel) were determined by RT-qPCR assays [Two-way ANOVA, *F*_*(3, 8)*_ = 18.60, *p*=0.0006 (Serum stimulation). *F* _*(3,8)*_ = 7.112, *p* = 0.012 (IFNγ sensitization); post-hoc Tukey’s HSD, *\*p* < 0.05, *\*\*p* < 0.01, *\*\*\*p* < 0.001, *\*\*\*\*p* < 0.001]. (F and G) Immunocytochemical analysis of PD-L1 in the TAP-EP400NL inducible Flp-In™ T-REx™ cell line. Cells were treated with either a combination of serum stimulation and tetracycline induction (F) or a combination of IFNγ sensitization and tetracycline induction (G) followed by the staining with the anti-PD-L1 and anti-EP400NL antibodies. Nuclei were counterstained with 4,6-diamidino-2-phenylindole (DAPI). Scale bar denotes 5 µm.

IFNγ also increased PD-L1 mRNA expression by approximately 2-fold and additional treatment of either serum stimulation or tetracycline induction resulted in an additive effect (Figure 6C). Interestingly, PD-L1 gene expression was synergistically upregulated up to eight-fold by triple treatments with IFNγ sensitization, serum stimulation, and tetracycline induction (IFNγ+/Serum+/Tetra+) (Figure 6C, strapped bar). A similar pattern of PD-L1 protein expression was observed in the triple-treated cells (IFNγ+/Serum+/Tetra+) with a 2.5-fold increase compared to the control cells (Figure 6D, anti-PD-L1 immunoblot).

To establish the specificity of EP400NL coactivator function in cMyc and IFNγ mediated PD-L1 expression, two deletion mutants (Δ246-260 and Δ360-419) that both lacking the coactivator function were examined for regulating PD-L1 expression in the presence of either serum stimulation or IFNγ sensitization. In contrast to wild type EP400NL, neither mutant induced expression of PD-L1 mRNA (Figure 6E) or protein (data not shown). These data confirm that the regions of EP400NL between amino acids 246-260 and 360-419 are important in regulating the EP400NL-mediated coactivator function of PD-L1 gene activation.

PD-L1 protein upregulation was also confirmed via confocal microscopy. Flp-In™ T-REx™ cells were tetracycline-induced to express EP400NL then stained with anti-PD-L1, anti-EP400NL, and DAPI after exposure to either serum stimulation or IFNγ sensitization. An increased expression of EP400NL was detected after tetracycline induction, but no alteration in PD-L1 expression was observed following serum-starvation or treatment without IFNγ sensitization regardless of tetracycline induction (Figure 6F and 6G, top two panels). On the contrary, all cells expressed a significantly higher level of PD-L1 after serum stimulation or IFNγ sensitization, which was further increased after EP400NL induction by tetracycline (Figure 6F and 6G, bottom two panels). Taken together, these results demonstrate that Myc and IFNγ require the presence of EP400NL to increase PD-L1 expression.

CRISPR/Cas9 mediated EP400NL indels were introduced to study the loss of EP400NL gene function in mediating Myc and IFNγ mediated PD-L1 expression. To confirm the presence of indels at the target sites, high-resolution melting peak analysis (HRM) was employed to examine melt peak shifting and relative fluorescent signal differences of the dissociation curves (Supplementary Figure 1). The result shows the alteration of genomic DNA at the intended deletion sites and significantly decreased protein expression of endogenous EP400NL from the two-biological replicated EP400NL indels cell lines (Figures 7A and 7B, anti-EP400NL immunoblots). Interestingly, the targeted deletion of EP400NL did not completely abrogate the endogenous EP400NL protein, suggesting the complex gene structure and alternative RNA splicing of EP400NL transcripts (UniProtKB - Q6ZTU2).

**Figure 7.**
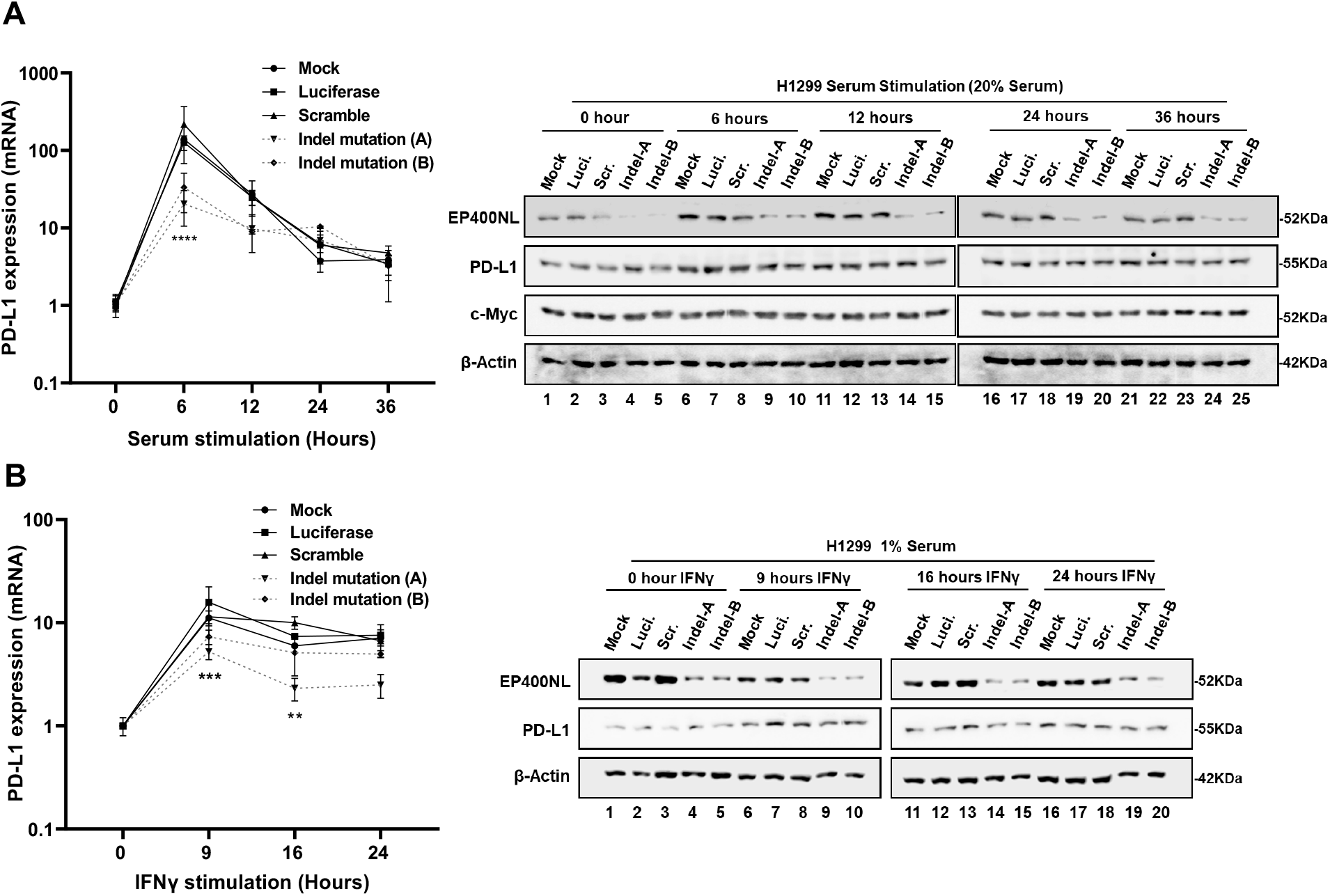
EP400NL enhances both Myc and IFNγ mediated PD-L1 expression. (A and B) PD-L1 mRNA expression levels in H1299 cell lines carrying EP400NL indels. These experiments were conducted under the condition of either serum stimulation (A) or IFNγ sensitization (B). The basal PD-L1 expression level of each cell line set to one and the expression levels of PD-L1 were calculated against their basal expression levels [Two-way ANOVA, *F*_*(4,25)*_ = 21.10, *p* < 0.0001 (Serum stimulation), *F*_*(3,20)*_ = 36.56, *p* < 0.0001 (IFNγ sensitization); post-hoc Tukey’s HSD, *\*\*p* < 0.001 (Scr. versus Indel-A, IFNγ sensitization), *\*\*\*p* < 0.001 (Luci. Vs. Indel-A, IFNγ sensitization) *\*\*\*\*p* < 0.0001 (Scr. Vs. Indel-A or Indel-B, serum stimulation)]. EP400NL immunoblots for H1299 cell lines established by CRISPR-Cas9 system (A, B, right panels): Mock, gRNA-luciferase (Luci.), gRNA-scrambled random sequence (Scr.), gRNA-EP400NL-A (Indel-A), and gRNA-EP400NL-B (Indel-B). Protein expression levels of endogenous PD-L1 under either serum stimulation (A, right panel) or IFNγ sensitization (B, right panel) were examined by immunoblots, respectively.

To investigate if the downregulation of EP400NL can affect Myc and IFNγ mediated PD-L1 expression, the two EP400NL indel lines (Indel-A, Indel-B) were used to examine PD-L1 expression in the time course of serum or IFNγ stimulation. Compared to H1299 control cells showing a dramatic increase of PD-L1 mRNA up to approximately 100 fold right after 6 hours of serum stimulation, polyclonal H1299 cells carrying EP400NL indels show compromised gene induction profiles that only gives about 30-fold of the transcript increase at the same time interval (Figure 7A, left panel). IFNγ sensitization shows decreased PD-L1 expression from the EP400NL indel cell lines, with a significantly lower expression level in one of the biological replicates (Indel-A) (Figure 7B, left panel). Taken together, the loss of function studies using CRISPR/Cas9-mediated indels confirmed the critical role of EP400NL in driving both Myc and IFNγ-mediated PD-L1 transcriptional activation. However, the weaker transcriptional activity of PD-L1 gene caused by EP400NL indels failed to show significant downregulation of PD-L1 protein levels (Figures 7A and 7B, anti-PD-L1 immunoblot).

## DISCUSSION

In this study, we found EP400NL, previously thought to be a pseudogene, forms a chromatin-remodeling complex similar to the hNuA4 histone acetyltransferase complex or EP400 complex. EP400NL is not an ATPase but our studies demonstrate that its nuclear multi-protein complex can catalyse an ATP-dependent H2A.Z deposition. In addition, upregulation of EP400NL enhances PD-L1 gene expression mediated by cMyc and IFNγ-regulated transcription factors.

EP400NL appears to form a nuclear complex similar to the TIP60-deficient EP400 complex but containing the BRG1 ATPase. The association of BRG1 with the EP400NL complex may be relatively weaker as the amount of BRG1 co-purified by EP400NL is minimal compared to other shared core subunits of the complex. The hNuA4 complex not only exhibits HAT activity but also catalyses histone acetylation-induced H2A.Z deposition (6,40,43,44). Specifically, the acetylated histone tails can stimulate and enhance the H2A.Z exchange reaction by the cooperative action of RuvBL1 and RuvBL2 via their ATPase activities (41,45). Since the synthetic chromatin in our H2A.Z deposition assay was prepared using recombinant core histones without pre-acetylation, it will be interesting to test if pre-acetylation of H2A on the chromatin can further enhance the EP400NL complex-mediated H2A.Z deposition activity.

Several identified proteins associated with EP400NL are also shared with the hNuA4 complex, including DMAP1, RuvBL1, RuvBL2, BAF53, MRGBP, and YEATS4 (2,46). It is well known that bromodomain specifically binds to acetylated lysine residues on histone tails, resulting in the targeted recruitment of the bromodomain-containing protein complexes towards the acetylated chromatin for the transcription regulation (47). BRD8 (Bromodomain-containing protein 8) is a transcriptional coactivator recruited by hormone nuclear receptors, in which isoforms of BRD8 were reported as the subunit of hNuA4 histone acetyltransferase complex for transcriptional activation (2,48). In addition, BRG1 contains a highly conserved bromodomain at the C-terminus, which might confer a missing ATPase function to the EP400NL complex (49). Given the similarity of the complex composition, it seems feasible in the cells that EP400 and EP400NL may be interchangeable in some specific circumstances of the chromatin remodelling process. The bromodomain-containing BRD8 and BRG1 of the EP400NL complex can be recruited by a DNA-binding transcription factor and the interaction would be significantly enhanced by the hyper-acetylated chromatin. Interestingly, the BRG1-containing BAF complex is also known to play a role in H2A.Z deposition in embryonic stem cells but little is understood about its molecular mechanism (50). It is tempting to speculate that the EP400NL complex is a carrier of BRG1 in certain genomic regions of H2A.Z deposition activity, however, the connection between BRG1 and EP400NL complex-mediated H2A.Z deposition and the coactivator function remains to be further elucidated.

Despite a higher protein amount of two EP400NL deletion mutants (Δ246-260, Δ360-419) in the immunoprecipitates, they showed a weaker interaction with BAF53 than the full-length EP400NL and lost the coactivator activity of Myc in the luciferase reporter assay (Figures 4B and 4C). Since BAF53 functions as a critical Myc-interacting nuclear cofactor for oncogenic transformation (51), these regions (246-260aa., 360-419aa.) of EP400NL would play a role in the association with BAF53 for co-activating Myc-mediated transcription.

Although cMyc level was not altered significantly by the serum stimulation in our EP400NL inducible Flp-In™ T-REx™ cells, serum stimulation in the presence of upregulated EP400NL appears to enhance the Myc binding to the PD-L1 promoter. ChIP data shows that the recruitment of EP400NL to the target promoter requires the binding of DNA binding transcription factors such as cMyc or other unknown serum-stimulated factors at the promoter (52). The co-immunoprecipitation studies of cMyc and EP400NL complex and their enrichment at the Myc binding site of PD-L1 gene support our model that cMyc recruits epigenetic modifying complexes to remodel the target gene promoters for transcriptional activation (Figure 8). Additionally, the PD-L1 expression pattern from cells with EP400NL indels suggests that EP400NL is required at an early stage of the transcriptional activation within 6 hours of serum stimulation (Figure 7A, left panel). PD-L1 overexpression is a predictor of recurrent cancer incidence and is associated with a poor prognosis of cancer patients (10,53). Consistent with the role in the PD-L1 gene regulation, cMyc expression correlates with PD-L1 expression in NSCLC (11). A previous study demonstrated that the tobacco-specific carcinogen NNK can trigger higher PD-L1 expression and cause primarily lung adenoma in mice (18,54). Moreover, a meta-analysis of mRNA expression profile identified EP400NL being upregulated in lung adenocarcinoma tissue from the cancer patients who have a smoking history (55). Multiple transcription factors and epigenetic protein complexes including the hNuA4, EP400, and BRG1-containing BAF complexes have been identified as interacting partners with Myc to induce and maintain cancerous phenotypes (2,7,8,29,32-34,51,56-59). Our studies show that EP400NL forms a unique nuclear complex with BRG1 and contributes to Myc-mediated PD-L1 gene expression, providing a potential therapeutic target to suppress cancer-associated PD-L1 gene expression.

**Figure 8.**
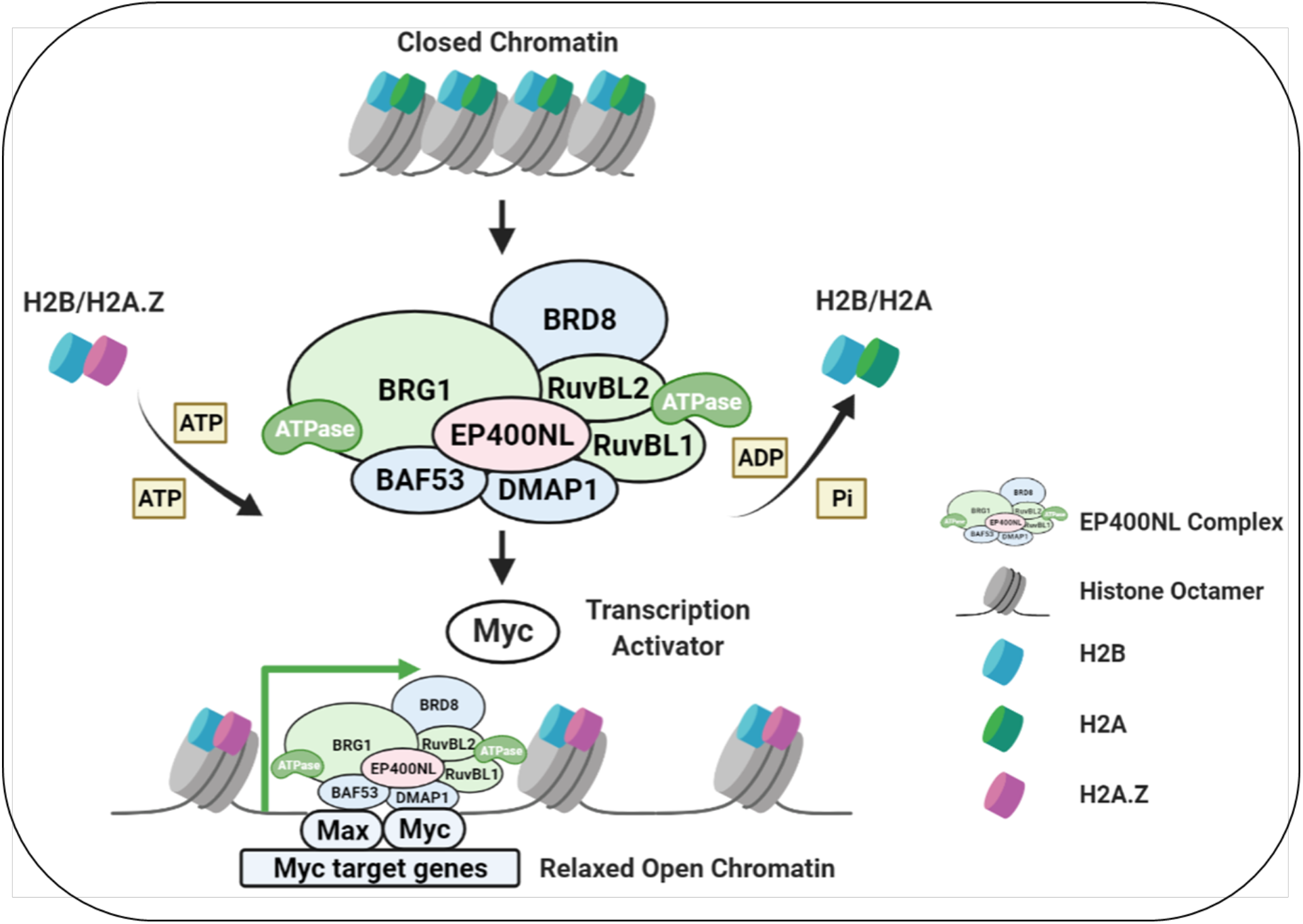
Working model of cMyc-dependent recruitment of EP400NL complex and transcriptional activation of target genes. EP400NL forms a unique chromatin remodeling complex that can transform the closed chromatins into open chromatins by its ATP-dependent H2A.Z deposition activity. The sequence-specific DNA binding transcription activator such as Myc recruits EP400NL complex at the target promoter and allows the localised chromatin remodelling for the transcriptional activation.

EP400NL is only about 13.4% the size of EP400 but with a high protein sequence similarity up to 92% specifically to the N-terminal domain of the EP400. Despite being annotated as an EP400 pseudogene 1 (EP400P1), not only can it be transcribed and translated but also seems to be functional. Our results suggest that the EP400NL complex exhibits H2A.Z deposition activity and the transcriptional coactivator function which are similar to the neighbouring paralog, EP400 complex, but with a stronger coactivator function possibly through interacting with BRG1. Gene expression profiling of 54 tissues from GTEx RNA-sequence shows that EP400 is ubiquitously expressed at a high level in many tissues whereas higher EP400NL expression is restricted only in the testis and pituitary gland, suggesting a specialised function in those tissues. The exon regions of EP400NL are highly conserved across mammals, suggesting that the unique function is conferred by EP400NL may not be redundant with EP400. Although EP400NL modifies Myc and IFNγ mediated PD-L1 expression by forming a unique nuclear complex, the specific functional domains and direct binding partners of EP400NL remain to be identified. Therefore, elucidating the direct binding domain of EP400NL and how these interacting candidate proteins are recruited into the complex will be a focus of future work. Furthermore, investigation of EP400NL-induced whole transcriptome changes and genome-wide ChIP-sequencing will provide better insight on how EP400NL contributes to the human epigenome maintenance and gene regulation especially in connection with Myc-dependent cellular proliferation and oncogenic transformation.

## DATA AVAILABILITY

Not applicable.

## SUPPLEMENTARY DATA

+ Supplementary Table1. Antibodies for western-blot analysis

+ Supplementary Table 2. Primers for ChIP q-PCR assay

+ Supplementary Figure 1. Melt peak shifting and relative fluorescent signal differences of the dissociation curves

## ACKNOWLEDGEMENT

We thank Mr. Trevor Loo for his excellent technical support for mass spectrometric analysis. We acknowledge the Manawatu Microscopy and Imaging Centre for confocal microscopy use and their excellent technical assistance.

## REFERENCES

1. Doyon, Y., Selleck, W., Lane, W.S., Tan, S. and Cote, J. (2004) Structural and functional conservation of the NuA4 histone acetyltransferase complex from yeast to humans. Mol Cell Biol, 24, 1884–1896.

2. Yamada, H.Y. (2012), Colorectal Cancer Biology-From Genes to Tumor. Intechopen.

3. Lu, P.Y.T., Levesque, N. and Kobor, M.S. (2009) NuA4 and SWR1-C: two chromatin-modifying complexes with overlapping functions and components. Biochemistry and Cell Biology, 87, 799–815.

4. Xu, Y., Ayrapetov, M.K., Xu, C., Gursoy-Yuzugullu, O., Hu, Y.D. and Price, B.D. (2012) Histone H2A.Z Controls a Critical Chromatin Remodeling Step Required for DNA Double-Strand Break Repair. Molecular Cell, 48, 723–733.

5. Pradhan, S.K., Su, T., Yen, L.D., Jacquet, K., Cote, J., Kurdistani, S. and Carey, M. (2016) Chromatin Remodeler EP400 Deposits H3.3 into Promoters and Enhancers During Gene Activation. Faseb J, 30.

6. Giaimo, B.D., Ferrante, F., Herchenrother, A., Hake, S.B. and Borggrefe, T. (2019) The histone variant H2A.Z in gene regulation. Epigenet Chromatin, 12.

7. Tworkowski, K.A., Chakraborty, A.A., Samuelson, A.V., Seger, Y.R., Narita, M., Hannon, G.J., Lowe, S.W. and Tansey, W.P. (2008) Adenovirus E1A targets p400 to induce the cellular oncoprotein Myc. Proc Natl Acad Sci U S A, 105, 6103–6108.

8. Zhao, L.J., Loewenstein, P.M. and Green, M. (2017) Enhanced MYC association with the NuA4 histone acetyltransferase complex mediated by the adenovirus E1A N-terminal domain activates a subset of MYC target genes highly expressed in cancer cells. Genes Cancer, 8, 752–761.

9. Jiang, X., Wang, J., Deng, X., Xiong, F., Ge, J., Xiang, B., Wu, X., Ma, J., Zhou, M., Li, X. et al. (2019) Role of the tumor microenvironment in PD-L1/PD-1-mediated tumor immune escape. Mol Cancer, 18, 10.

10. Casey, S.C., Tong, L., Li, Y.L., Do, R., Walz, S., Fitzgerald, K.N., Gouw, A.M., Baylot, V., Gutgemann, I., Eilers, M. et al. (2016) MYC regulates the antitumor immune response through CD47 and PD-L1. Science, 352, 227–231.

11. Kim, E.Y., Kim, A., Kim, S.K. and Chang, Y.S. (2017) MYC expression correlates with PD-L1 expression in non-small cell lung cancer. Lung Cancer, 110, 63–67.

12. Karachaliou, N., Gonzalez-Cao, M., Crespo, G., Drozdowskyj, A., Aldeguer, E., Gimenez-Capitan, A., Teixido, C., Molina-Vila, M.A., Viteri, S., Gil, M.D. et al. (2018) Interferon gamma, an important marker of response to immune checkpoint blockade in non-small cell lung cancer and melanoma patients. Ther Adv Med Oncol, 10.

13. Seliger, B. (2019) Basis of PD1/PD-L1 Therapies. J Clin Med, 8.

14. Ju, X.L., Zhang, H., Zhou, Z.D. and Wang, Q. (2020) Regulation of PD-L1 expression in cancer and clinical implications in immunotherapy. Am J Cancer Res, 10, 1–11.

15. Keir, M.E., Butte, M.J., Freeman, G.J. and Sharpel, A.H. (2008) PD-1 and its ligands in tolerance and immunity. Annu Rev Immunol, 26, 677–704.

16. Chen, J., Jiang, C.C., Jin, L. and Zhang, X.D. (2016) Regulation of PD-L1: a novel role of prosurvival signalling in cancer. Ann Oncol, 27, 409–416.

17. Jiang, X.J., Wang, J., Deng, X.Y., Xiong, F., Ge, J.S., Xiang, B., Wu, X., Ma, J., Zhou, M., Li, X.L. et al. (2019) Role of the tumor microenvironment in PD-L1/PD-1-mediated tumor immune escape. Molecular Cancer, 18.

18. Lastwika, K.J., Wilson, W., Li, Q.K., Norris, J., Xu, H.Y., Ghazarian, S.R., Kitagawa, H., Kawabata, S., Taube, J.M., Yao, S. et al. (2016) Control of PD-L1 Expression by Oncogenic Activation of the AKT-mTOR Pathway in Non-Small Cell Lung Cancer. Cancer Research, 76, 227–238.

19. Moon, J.W., Kong, S.K., Kim, B.S., Kim, H.J., Lim, H., Noh, K., Kim, Y., Choi, J.W., Lee, J.H. and Kim, Y.S. (2017) IFN gamma induces PD-L1 overexpression by JAK2/STAT1/IRF-1 signaling in EBV-positive gastric carcinoma. Sci Rep-Uk, 7.

20. Gowrishankar, K., Gunatilake, D., Gallagher, S., Tiffen, J. and Hersey, P. (2014) Regulation of PD-L1 expression in human melanoma by NF-kappa B. Cancer Research, 74.

21. Gong, J., Chehrazi-Raffle, A., Reddi, S. and Salgia, R. (2018) Development of PD-1 and PD-L1 inhibitors as a form of cancer immunotherapy: a comprehensive review of registration trials and future considerations. J Immunother Cancer, 6.

22. Sun, C., Mezzadra, R. and Schumacher, T.N. (2018) Regulation and Function of the PD-L1 Checkpoint. Immunity, 48, 434–452.

23. Garcia-Diaz, A., Shin, D.S., Moreno, B.H., Saco, J., Escuin-Ordinas, H., Rodriguez, G.A., Zaretsky, J.M., Sun, L., Hugo, W., Wang, X.Y. et al. (2019) Interferon Receptor Signaling Pathways Regulating PD-L1 and PD-L2 Expression (vol 19, pg 1189, 2017). Cell Rep, 29, 3766–3766.

24. Antonangeli, F., Natalini, A., Garassino, M.C., Sica, A., Santoni, A. and Di Rosa, F. (2020) Regulation of PD-L1 Expression by NF-kappa B in Cancer. Front Immunol, 11.

25. Smith, R.J., Savoian, M.S., Weber, L.E. and Park, J.H. (2016) Ataxia telangiectasia mutated (ATM) interacts with p400 ATPase for an efficient DNA damage response. Bmc Molecular Biology, 17.

26. Slaymaker, I.M., Gao, L.Y., Zetsche, B., Scott, D.A., Yan, W.X. and Zhang, F. (2016) Rationally engineered Cas9 nucleases with improved specificity. Science, 351, 84–88.

27. Ghosh, K., Tang, M., Kumari, N., Nandy, A., Basu, S., Mall, D.P., Rai, K. and Biswas, D. (2018) Positive Regulation of Transcription by Human ZMYND8 through Its Association with P-TEFb Complex. Cell Rep, 24, 2141-+.

28. Kufe, D.W., Ma, US). (2020) United States.

29. Fuchs, M., Gerber, J., Drapkin, R., Sif, S., Ikura, T., Ogryzko, V., Lane, W.S., Nakatani, Y. and Livingston, D.M. (2001) The p400 complex is an essential E1A transformation target. Cell, 106, 297–307.

30. Stielow, B., Sapetschnig, A., Wink, C., Kruger, I. and Suske, G. (2008) SUMO-modified Sp3 represses transcription by provoking local heterochromatic gene silencing. Embo Rep, 9, 899–906.

31. Haas, J., Bloesel, D., Bacher, S., Kracht, M. and Schmitz, M.L. (2020) Chromatin Targeting of HIPK2 Leads to Acetylation-Dependent Chromatin Decondensation. Front Cell Dev Biol, 8.

32. McMahon, S.B., Van Buskirk, H.A., Dugan, K.A., Copeland, T.D. and Cole, M.D. (1998) The novel ATM-related protein TRRAP is an essential cofactor for the c-Myc and E2F oncoproteins. Cell, 94, 363–374.

33. Nikiforov, M.A., Chandriani, S., Park, J., Kotenko, I., Matheos, D., Johnsson, A., McMahon, S.B. and Cole, M.D. (2002) TRRAP-dependent and TRRAP-independent transcriptional activation by Myc family oncoproteins. Mol Cell Biol, 22, 5054–5063.

34. Liu, X.H., Tesfai, J., Evrard, Y.A., Dent, S.Y.R. and Martinez, E. (2003) C-Myc transformation domain recruits the human STAGA complex and requires TRRAP and GCN5 acetylase activity for transcription activation. Journal of Biological Chemistry, 278, 20405–20412.

35. Baek, H.J., Malik, S., Qin, J. and Roeder, R.G. (2002) Requirement of TRAP/mediator for both activator-independent and activator-dependent transcription in conjunction with TFIID-associated TAF(II)s. Molecular and Cellular Biology, 22, 2842–2852.

36. Muchir, A., Medioni, J., Laluc, M., Massart, C., Arimura, T., Van Der Kooi, A.J., Desguerre, I., Mayer, M., Ferrer, X., Briault, S. et al. (2004) Nuclear envelope alterations in fibroblasts from patients with muscular dystrophy, cardiomyopathy, and partial lipodystrophy carrying lamin A/C gene mutations. Muscle Nerve, 30, 444–450.

37. Bruce, D.L., Macartney, T., Yong, W., Shou, W. and Sapkota, G.P. (2012) Protein phosphatase 5 modulates SMAD3 function in the transforming growth factor-beta pathway. Cell Signal, 24, 1999–2006.

38. Doyon, Y. and Cote, J. (2004) The highly conserved and multifunctional NuA4 HAT complex. Curr Opin Genet Dev, 14, 147–154.

39. Lopez-Perrote, A., Alatwi, H.E., Torreira, E., Ismail, A., Ayora, S., Downs, J.A. and Llorca, O. (2014) Structure of Yin Yang 1 Oligomers That Cooperate with RuvBL1-RuvBL2 ATPases. Journal of Biological Chemistry, 289, 22614–22629.

40. Mizuguchi, G., Shen, X., Landry, J., Wu, W.H., Sen, S. and Wu, C. (2004) ATP-driven exchange of histone H2AZ variant catalyzed by SWR1 chromatin remodeling complex. Science, 303, 343–348.

41. Puri, T., Wendler, P., Sigala, B., Saibil, H. and Tsaneva, I.R. (2007) Dodecameric structure and ATPase activity of the human TIP48/TIP49 complex. J Mol Biol, 366, 179–192.

42. Wu, Q., Lian, J.B., Stein, J.L., Stein, G.S., Nickerson, J.A. and Imbalzano, A.N. (2017) The BRG1 ATPase of human SWI/SNF chromatin remodeling enzymes as a driver of cancer. Epigenomics-Uk, 9, 919–931.

43. Altaf, M., Auger, A., Monnet-Saksouk, J., Brodeur, J., Piquet, S., Cramet, M., Bouchard, N., Lacoste, N., Utley, R.T., Gaudreau, L. et al. (2010) NuA4-dependent Acetylation of Nucleosomal Histones H4 and H2A Directly Stimulates Incorporation of H2A.Z by the SWR1 Complex. Journal of Biological Chemistry, 285, 15966–15977.

44. Ranjan, A., Mizuguchi, G., FitzGerald, P.C., Wei, D., Wang, F., Huang, Y.Z., Luk, E., Woodcock, C.L. and Wu, C. (2013) Nucleosome-free Region Dominates Histone Acetylation in Targeting SWR1 to Promoters for H2A.Z Replacement. Cell, 154, 1232–1245.

45. Choi, J., Heo, K. and An, W.J. (2009) Cooperative action of TIP48 and TIP49 in H2A.Z exchange catalyzed by acetylation of nucleosomal H2A. Nucleic Acids Res, 37, 5993–6007.

46. Doyon, Y. and Cote, J. (2004) The highly conserved and multifunctional NuA4 HAT complex. Current Opinion in Genetics & Development, 14, 147–154.

47. Fujisawa, T. and Filippakopoulos, P. (2017) Functions of bromodomain-containing proteins and their roles in homeostasis and cancer. Nat Rev Mol Cell Bio, 18, 246–262.

48. Cai, Y., Jin, J., Tomomori-Sato, C., Sato, S., Sorokina, I., Parmely, T.J., Conaway, R.C. and Conaway, J.W. (2003) Identification of new subunits of the multiprotein mammalian TRRAP/TIP60-containing histone acetyltransferase complex. J Biol Chem, 278, 42733–42736.

49. Sanchez, J.C., Zhang, L.Y., Evoli, S., Schnicker, N.J., Nunez-Hernandez, M., Yu, L.P., Wereszczynski, J., Pufall, M.A. and Musselman, C.A. (2020) The molecular basis of selective DNA binding by the BRG1 AT-hook and bromodomain. Bba-Gene Regul Mech, 1863.

50. Hainer, S.J. and Fazzio, T.G. (2015) Regulation of Nucleosome Architecture and Factor Binding Revealed by Nuclease Footprinting of the ESC Genome. Cell Rep, 13, 61–69.

51. Park, J., Wood, M.A. and Cole, M.D. (2002) BAF53 forms distinct nuclear complexes and functions as a critical c-Myc-interacting nuclear cofactor for oncogenic transformation. Molecular and Cellular Biology, 22, 1307–1316.

52. Wang, Z., Wang, P., Li, Y., Peng, H., Zhu, Y., Mohandas, N. and Liu, J. (2021) Interplay between cofactors and transcription factors in hematopoiesis and hematological malignancies. Signal Transduct Target Ther, 6, 24.

53. Wang, X., Teng, F.F., Kong, L. and Yu, J.M. (2016) PD-L1 expression in human cancers and its association with clinical outcomes. Oncotargets Ther, 9, 5023–5039.

54. West, K.A., Linnoila, I.R., Belinsky, S.A., Harris, C.C. and Dennis, P.A. (2004) Tobacco carcinogen-induced cellular transformation increases activation of the phosphatidylinositol 3 ‘-Kinase/Akt pathway in vitro and in vivo. Cancer Research, 64, 446–451.

55. He, X.N., Zhang, C., Shi, C. and Lu, Q.Q. (2018) Meta-analysis of mRNA expression profiles to identify differentially expressed genes in lung adenocarcinoma tissue from smokers and non-smokers. Oncol Rep, 39, 929–938.

56. Park, J., Kunjibettu, S., McMahon, S.B. and Cole, M.D. (2001) The ATM-related domain of TRRAP is required for histone acetyltransferase recruitment and Myc-dependent oncogenesis. Gene Dev, 15, 1619–1624.

57. Frank, S.R., Parisi, T., Taubert, S., Fernandez, P., Fuchs, M., Chan, H.M., Livingston, D.M. and Amati, B. (2003) MYC recruits the TIP60 histone acetyltransferase complex to chromatin. Embo Rep, 4, 575–580.

58. Cowling, V.H. and Cole, M.D. (2006) Mechanism of transcriptional activation by the Myc oncoproteins. Semin Cancer Biol, 16, 242–252.

59. Vita, M. and Henriksson, M. (2006) The Myc oncoprotein as a therapeutic target for human cancer. Semin Cancer Biol, 16, 318–330.

