## Supplementary Data for "EP400NL is required for cMyc-mediated PD-L1 gene activation by forming a transcriptional coactivator complex"

### Supplementary Tables and Figures

**Zidong (Andy) Li**

**26.05.2021**

**Supplementary table 1. Antibodies for western-blot analysis**

| Antibodies | Dilution in TBST | Information |
|---|---|---|
| Anti-EP400NL | 1:1000 | HPA068417, Sigma-Aldrich |
| Anti-CBP | 1:500 | Sc-32998, Santa-Cruz Biotechnology |
| Anti-FLAG | 1:1000 | A8592, Sigma-Aldrich |
| Anti-BRG1 | 1:1000 | ab70558, Abcam, USA |
| Anti-BRD8 | 1:10000 | ab17969, Abcam, USA |
| Anti-BAF53 | 1:100 | A kind gift from Dr Cole |
| Anti-cMyc | 1:500 | MCA1929, Bio-Rad |
| Anti-PPM1B | 1:500 | AF4396, R&D System Minneapolis, USA |
| Anti-β-actin | 1:5000 | NB600-501, NOVUS, USA |
| Anti-Lamin A/C | 1:10000 | ab108595, Abcam, USA |
| Anti-TIP48 | 1:500 | A kind gift from Dr Cole |
| Anti-MED30 | 1:500 | MBS9609934, MYBioSource |
| Anti-TIP49 | 1:500 | A kind gift from Dr Cole |
| Anti-PD-L1 | 1:500 | #14-5982-82, Invitrogen |

**Supplementary table 2. Primers for ChIP q-PCR assay**

| | | |
|---|---|---|
| **P21 (Non-specific)** | Forward Primer | 5'-CACTGCAATTTGGCCCAGA-3' |
| | Reverse Primer | 5'-GTGCAGTAGAGAATTATTCCACATTTG-3' |
| **Gal4-DNA binding site** | Forward Primer | 5'-CTTATGGTACTGTAACTGAGCTAAC-3' |
| | Reverse Primer | 5'-GCGGGACTATGGTTGCTGAC-3' |
| **GAPDH (Non-specific)** | Forward Primer | 5'-TACTAGCGGTTTTACGGGCG-3' |
| | Reverse Primer | 5'-TCGAACAGGAGGAGCAGAGAGCGA-3' |
| **Promoter region of PD-L1** | Forward Primer | 5'-CATATGGGTCTGCTGCTGAC-3' |
| | Reverse Primer | 5'-CAACAAGCCAACATCTGAAC-3' |

**Supplementary Figure 1**

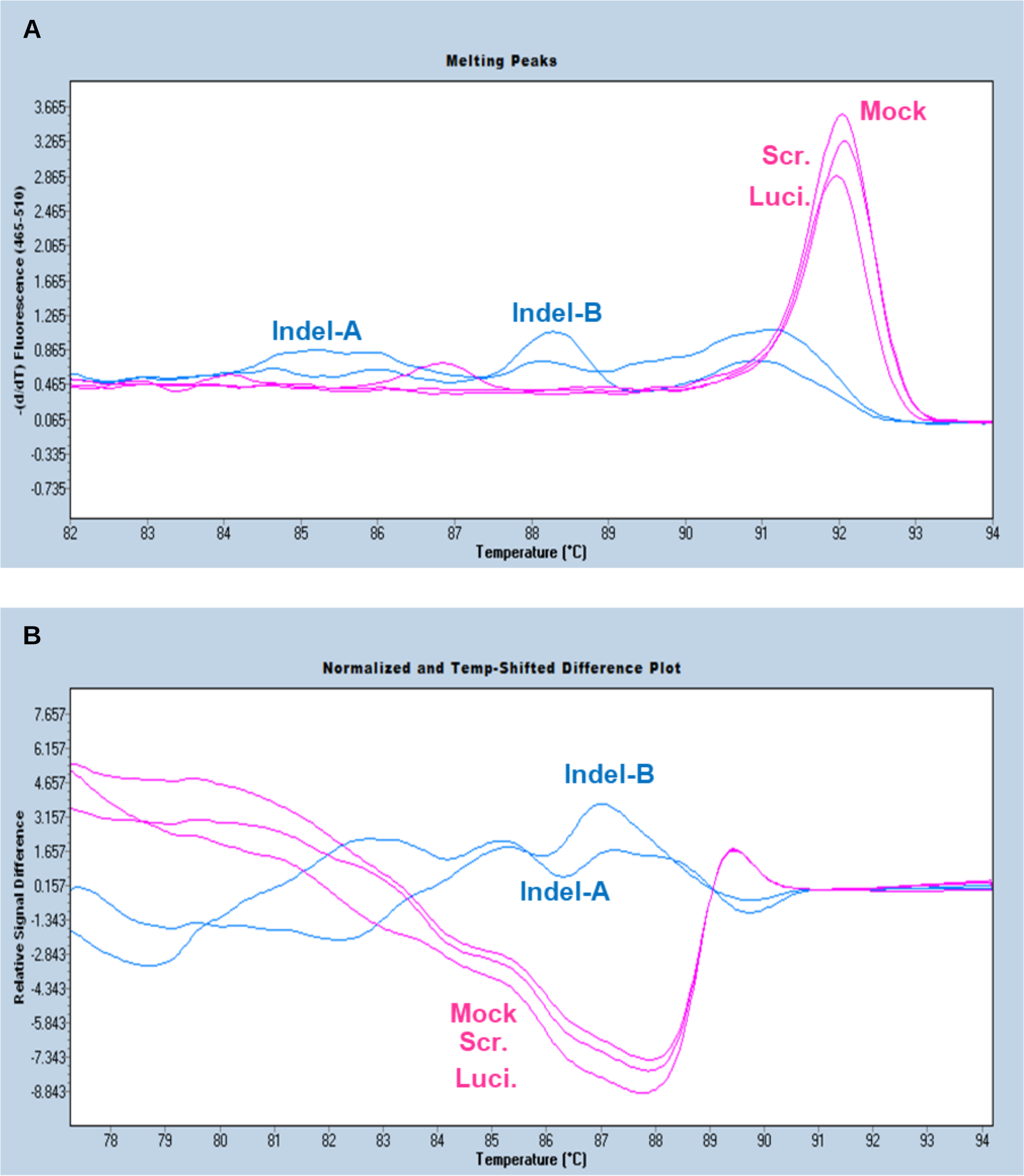


**Supplementary Figure 1. Melt peak shifting (A) and relative fluorescent signal differences of the dissociation curves (B)**

(A) Melting peak shifts were identified from the two-biological replicated EP400NL indels cell lines (Indel-A, Indel-B) compared to the three control cell lines which are Mock, Luciferase, and Scramble respectively. (B) Normalized and Temp-shifted difference plot of the three control cell lines and the two-biological replicated EP400NL indels cell lines. Mock, Wild type H1299; Luci., gRNA-luciferase; Scr., gRNA-scrambled random sequence; Indel-A, gRNA-EP400NL-A; Indel-B, gRNA-EP400NL-B.
